# The remarkable complexity of the brain microbiome in health and disease

**DOI:** 10.1101/2023.02.06.527297

**Authors:** Xinyue Hu, Chris-Anne Mckenzie, Colin Smith, Juergen G. Haas, Richard Lathe

## Abstract

Microbes in human brain and their potential contribution to neurodegenerative conditions such as Alzheimer’s disease (AD) have long been debated. We recently developed a new method (the electronic tree of life, eToL) based on small subunit ribosomal RNA (rRNA) probes, further confirmed by large subunit rRNA analysis, to comprehensively address the spectrum of microorganisms in control and AD brain. We report a remarkable diversity of brain microbes in control brain. The most abundant are fungi, bacteria, and chloroplastida, and we report detailed identification of representative microbial species. The pattern is substantially conserved across different bilateran species from *Drosophila* to human. In terms of diversity, the human brain microbiome appears to be a subset (∼20%) of the gut microbiome. Adenovirus type C was the major virus found in human brain; other viruses were not well represented. However, the spectrum of brain microbes differed between individuals as well as between brain regions examined from single individuals (amygdala, cingulate cortex, hippocampus, hypothalamus); of these four regions, the highest microbial burden was in cingulate cortex. There was evidence of spreading of pathogens between brain regions in single individuals. Some microbes are over-represented in AD brain according to two measures: (i) absolute number of microbes normalized to endogenous human transcripts, and (ii) the number of brain specimens showing overabundance versus control. Species over-represented in AD brain according to both measures notably include bacteria (*Streptococcus, Staphylococcus*/*Bacillus, Sphingomonas*/*Ralstonia*) and fungi (*Acrocalymma*/*Altenaria*/*Aureobasidium* of the *Aspergillus* group; *Komagataella* of the *Candida* group, C*ortinarius* of the *Schizophyllum* group, and *Tausonia* of the *Cryptococcus* group), that are all related to known human pathogens. In addition, an uncharacterized chloroplastida (algae-related) species was more abundant in AD brain samples. Although these findings point to diverse microbial species, indicative of multiple causation, similar absolute levels of bacteria and fungi in AD brain samples could suggest synergy between pathogens. However, it is important to stress that not all AD samples were positive for these microbes, but this could be because the affected brain region(s) was not examined. These findings support the contention that infection, perhaps associated with declining immunity with age, may contribute to AD development.

## INTRODUCTION

Higher organisms evolved in an environment dominated by microbes, and tissues of higher organisms including plants are widely associated with diverse microbial populations (the ‘microbiome’). These ‘endozooites’ (Lathe & St Clair 2020) include bacteria, fungi, and other cellular organisms, as well as viruses that target microbial or host cells. Some microbes can have beneficial effects – illustrated by the symbiotic associations of mycorrhizal fungi and nitrogen-fixing bacteria with plant roots, and of ‘probiotic’ bacteria with the human gut – whereas others are pathogens that must be kept in check by the immune system. However, there is increasing recognition that tissues in higher organisms outside the gut harbor their own microbiomes, and extensive work has been carried out to characterize the microbial populations of skin and internal spaces including the lung, mouth, stomach, vagina, bladder, and the nasal mucosa, as well as blood and lymph ((Turnbaugh et al. 2007; Human Microbiome Project Consortium 2012); https://hmpdacc.org/). In addition to the gut and epithelial surfaces, there is evidence that endozoites are present within solid organs such as liver and kidney (I tthitaetrakool *et al*. 2016; Whiteside *et al*. 2015), raising the question of whether the brain houses its own microbiome (Lathe & St Clair 2020; Link 2021).

In recent decades studies on the brain microbiome have principally employed PCR methods, in some cases backed up by *in situ* techniques, and researchers have reported finding both bacteria and fungi in normal human brain samples (e.g., (Emery *et al*. 2017; Pisa *et al*. 2015)). However, PCR techniques only amplify sequences that correspond to the primers employed, and other researchers have sought to use next-generation sequencing and/or metagenomics, as illustrated by the finding of herpesviruses in brain tissue using a *k*-mer method (Readhead *et al*. 2018) and of fungi in cerebrospinal fluid (CSF) samples from patients with menigitis/encephalitis of unknown origin using metagenomics (Wilson *et al*. 2018; Wilson *et al*. 2019). Nevertheless, these findings have not been immune from challenge (Chorlton 2020), and, despite many impressive studies, none to date has addressed the entire microbiome, and few if any have addressed the absolute abundance of microbes in brain. Moreover, metagenomics is very demanding on computer processing and data storage. We therefore developed a different approach that is both less resource-intensive and neutral with regard to the identity of the species detected. The electronic tree of life (eToL) method is based on a net of over 1000 64-mer probes designed from the 16S/18S rRNA sequences of all known branches of life, including archaea, bacteria, chloroplastida, amoeobozoa, basal eukaryota, fungi, and metazoa; the technical details of this method were reported previously (Hu *et al*. 2022). We now apply this methodology to the study of the human brain microbiome in health and disease.

Since the time of Alzheimer and Fischer there has been debate about the possibility that select microbes might contribute to the neuroinflammation and neuronal damage that are seen in age-related diseases such as Alzheimer’s disease (AD) (Fischer 1910), a devastating neurodegenerative condition principally affecting the elderly (reviewed in (Masters *et al*. 2015)). This contention has been further fuelled by the finding that the signature protein of AD brain, amyloid β, is an antimicrobial peptide ((Soscia *et al*. 2010; Kumar *et al*. 2016), reviewed in (Moir *et al*. 2018)), and a similar function has now been attributed to the other protein deposited in AD brain, tau (Eimer *et al*. 2026), which suggests that the role of these deposits is to defend the brain against infection.

Moreover, apoliprotein E, the product of the *APOE* gene (a key determinant of AD risk) interacts directly with amyloid β and is thought to play a prominent antimicrobial role (e.g., (Petruk *et al*. 2021)). Indeed, immune decline with aging (immunosenescence) might predispose to infections of the brain, culminating in neurodegeneration (reviewed in (Lathe & St Clair 2023)). Diverse microbes including bacteria, yeasts, and viruses have been reported in AD brain (MacDonald 1986; Miklossy 2011; Branton *et al*. 2013; Itzhaki 2014; Pisa *et al*. 2015; Emery *et al*. 2017; Balin *et al*. 2018; Moné *et al*. 2023). Although no unique ‘AD microbe’ has so far been identified, these findings have prompted the speculation that microbial infection of the brain could contribute to AD development (Itzhaki *et al*. 2016).

However, these studies have without exception addressed a restricted range of species. We therefore used the eToL methodology to evaluate the full microbial spectrum in normal and AD brain, and to determine their absolute abundances, and have examined whether any specific microbes are differentially represented in AD brain. A preliminary report of this work has appeared (Hu *et al*. 2022), and we now present the full analysis.

## MATERIALS AND METHODS

### Brain RNA-seq datasets

The ‘Miami’ (Magistri *et al*. 2015) and ‘Rockefeller’ (Scheckel *et al*. 2016) human brain RNA-seq datasets are freely available online at the National Center for Biotechnology Information (NCBI). Mount Sinai Brain Bank (MSBB) RNA-seq datasets (Wang *et al*. 2018) were accessed, with permission, through the AD Knowledge Portal (https://adknowledgeportal.synapse.org/) sponsored by the National Institute on Aging (NIA), in compliance with Creative Commons Attribution 4.0 International Public License. The results published here are therefore in part based on data obtained from the AD Knowledge Portal (https://adknowledgeportal.org/). These data were generated from postmortem brain tissue collected through the Mount Sinai VA Medical Center Brain Bank and were provided by Dr Eric Schadt from Mount Sinai School of Medicine. The datasets were reconstructed (courtesy of B. Readhead) to make them comparable to the datasets analyzed in (Readhead *et al*. 2018). The multi-species, gut, and human and macaque brain with age RNA-seq datasets used in this work are also publicly available at NCBI. Sequence read archive (SRA) reference numbers for all datasets are listed in Table S1 in the supplementary material online. Because age and sex data were not available for some datasets, analysis of these factors was not possible.

### Edinburgh Brain Bank (EBB) neuropathology and deep sequencing

Frozen brain samples (four brain regions per individual: amygdala, cingulate cortex, hippocampus, and hypothalamus) from three control brains and six AD brains were obtained from the EBB. Analysis received ethical approval under generic Medical Research Council (MRC) ethical approval relating to EBB samples and were in compliance with the Human Tissue Act (Scotland) 2006 and subsequent legislation. The individuals in the AD category were diagnosed as ‘Alzheimer disease’ during their lifetimes, but postmortem analysis revealed that two individuals had some features of vascular dementia (VaD) or Lewy body dementia (LBD). However, because this is a common issue in the field, these individuals were retained within the broader ‘AD’ category. Because the integrity of one frozen sample was compromised, the EBB dataset was based on a total of 35 brain samples (summarized in **Table S2**).

#### Analysis of pathology

At post-mortem, fresh tissue samples (ca 3 × 2 × 1 cm) were taken and stored on ice packs until return to the laboratory. Using a Category 2 hood, fresh tissue samples were further dissected into 1 × 1 × 1cm cubes. Samples were frozen in liquid nitrogen vapour for 20 minutes before being transferred to a ™80 C freezer for long-term storage. All cases have detailed neuropathological examination using formalin-fixed paraffin embedded sections (FFPE). Routine analysis was done by hematoxylin and eosin (H&E) staining of 6 μm sections. Immunohistochemistry to assess neurodegenerative proteins was performed on 4 μm sections; tau analysis (no pretreatment, antibody AT8, Thermofisher MN1020, 1:2000, 30 minutes) and β-amyloid analysis (pretreatment with formic acid, antibody 4G8, DAKO, M087201-2, 1:100, 30 minutes). Tau pathology was assessed using Braak criteria (Braak *et al*. 2006), and amyloid pathology using Thal phase (Thal *et al*. 2002). Formal neuropathological assessment of AD employed National Institute of Aging (NIA-AA) criteria (Hyman *et al*. 2012).

#### Deep sequencing

Total RNA was extracted from frozen tissue using Qiagen RNeasy Plus Universal mini kit following the manufacturer’s instructions (Qiagen, Hilden, Germany). RNA samples were quantified using a Qubit 2.0 Fluorometer (ThermoFisher Scientific, Waltham, MA, USA) and RNA integrity was checked with the 4200 TapeStation (Agilent Technologies, Palo Alto, CA, USA). Human rRNA depletion was performed using the QIASeq FastSelect kit (Qiagen, Hilden, Germany). RNA sequencing library preparation used NEBNext Ultra II RNA Library Prep Kit for Illumina following the manufacturer’s recommendations (New England Biolabs, Ipswich, MA, USA). Briefly, enriched RNAs were fragmented for 15 minutes at 94°C. First and second strand cDNA was synthesized, cDNA fragments were end-repaired and universal adapters were ligated to the cDNA fragments, followed by limited-cycle PCR. Sequencing libraries were validated using the Agilent Tapestation 4200 (Agilent Technologies, Palo Alto, CA, USA), and quantified by using Qubit 2.0 Fluorometer (Invitrogen, Carlsbad, CA) as well as by quantitative PCR (Applied Biosystems, Carlsbad, CA, USA). The sequencing libraries were multiplexed and clustered on the flowcell. After clustering, the flowcell was loaded onto the Illumina NovaSeq 6000 instrument according to the manufacturer’s instructions. The samples were sequenced using a 2 × 150 paired-end (PE) configuration. Raw sequence data (.bcl files) generated from Illumina NovaSeq were converted into fastq files and demultiplexed using Illumina bcl2fastq program version 2.20. The datasets are publicly available at NCBI (**Table S1**)

### Probing for microbial, retroelement, and viral sequences

The eToL method is based on a net of >1000 64-mer probes designed from 16S/18S rRNA sequences of cellular organisms identified from the Open Tree of Life resource (https://opentreeoflife.org) extended to include bacteria according to the framework established by Schulz and colleagues (Schulz *et al*. 2017). The technical details of eToL have been reported previously (Hu *et al*. 2022). The workflow involves (i) generation of random probes from small ribosomal subunit sequences, (ii) sequence searching against the human transcriptome/genome using BLAST (basic local alignment search tool) (Altschul *et al*. 1990) and removal of all probes with significant matches to human sequences, generating a net of 1017 probes, (iii) use of the microbe-specific probe collection to retrieve non-human sequences from RNA-seq data (BLAST search), (iv) refiltering to remove any remaining human sequences, (v) normalization of readcounts to endogenous human housekeeping genes to give the number of microbial transcripts per host cell, (vi) species identification by contig assembly and databaset searching. BLAST searching was performed at the Edinburgh Compute and Data Facility (ECDF) Linux Compute Cluster (http://www.ecdf.ed.ac.uk/) (‘EDDIE’) at the University of Edinburgh. Other runs were performed online at NCBI (https://blast.ncbi.nlm.nih.gov/Blast.cgi). The extended eToL probelist also includes sequences corresponding to endogenous retroelements (long and short interspersed nuclear elements, SINEs/LINEs; and endogenous human retroviruses, HERVs) (Hu *et al*. 2022), and the same method was used to determine the abundances of different retroelement transcripts in RNA-seq data from different tissues.

The following (simplified) categories are employed by eToL: A, archaea; B, bacteria; C, chloroplastida (algae and plants), D, amoebozoa; E0, basal eukaryota (that may constitute a clade of their own); F, fungi; and H, holozoa/metazoa. Group G was not allocated. Microbial profiles were plotted using Morpheus software at the Broad Institute of MIT and Harvard (Cambridge, MA, USA; https://software.broadinstitute.org/morpheus/) either with or without conversion to log_2_. Cutoff values (specified in the figures) for cellular species of 3–5 reads per host cell were applied in most cases to exclude low-level (potentially contaminant) signals. For cellular microbes of interest, their presence was confirmed by downloading large subunit (23S/28S) rRNA, devising new probes according to the eToL method, and reprobing the RNA-seq data. As before, matches were retrieved, filtered against human sequences, and their identities determined by sequence searching on NCBI databases.

A variant approach was used for viruses. In previous work, Readhead and colleagues performed extensive analysis of viral sequences in human brain using a *k*-mer method (Readhead *et al*. 2018). Although potentially liable to false positives (Chorlton 2020), it is very unlikely to generate false negatives. We therefore took the top 20 viral groups identified by Readhead *et al*. that represent >99% of all viruses in their human brain samples analyzed. Because several of these viruses have internal regions of homology with the human genome (Hu *et al*. 2022), the complete genomes of all 20 viruses were ‘stripped’ to remove regions similar to human sequences. The complete stripped genomes were then used as probes for BLAST searching of RNA-seq libraries. Normalization to housekeeping gene expression was as for cellular sequences. A lower cutoff (0.03 reads per host cell) was used for viral sequences.

## RESULTS

### Cellular microbes in brain

We report a remarkable complexity of cellular microbes in normal human brain, principally fungi, bacteria, and chloroplastida. In addition, we detected archaea, amoebozoa, basal eukaryota, and even holozoa/metazoa (**Figure 1**). To identify representative species, matching sequences were retrieved, contigs assembled, and species identified by reference to sequence databases. To confirm the presence and identity of these species, the corresponding 23S/28S sequences were used to reprobe the RNA-seq datasets, contigs were generated, and identification was performed as before. The identities of representative microbial species determined by both 16S/18S and 23S/28S rRNA analysis are listed in **Table S3**.

**Figure 1.**
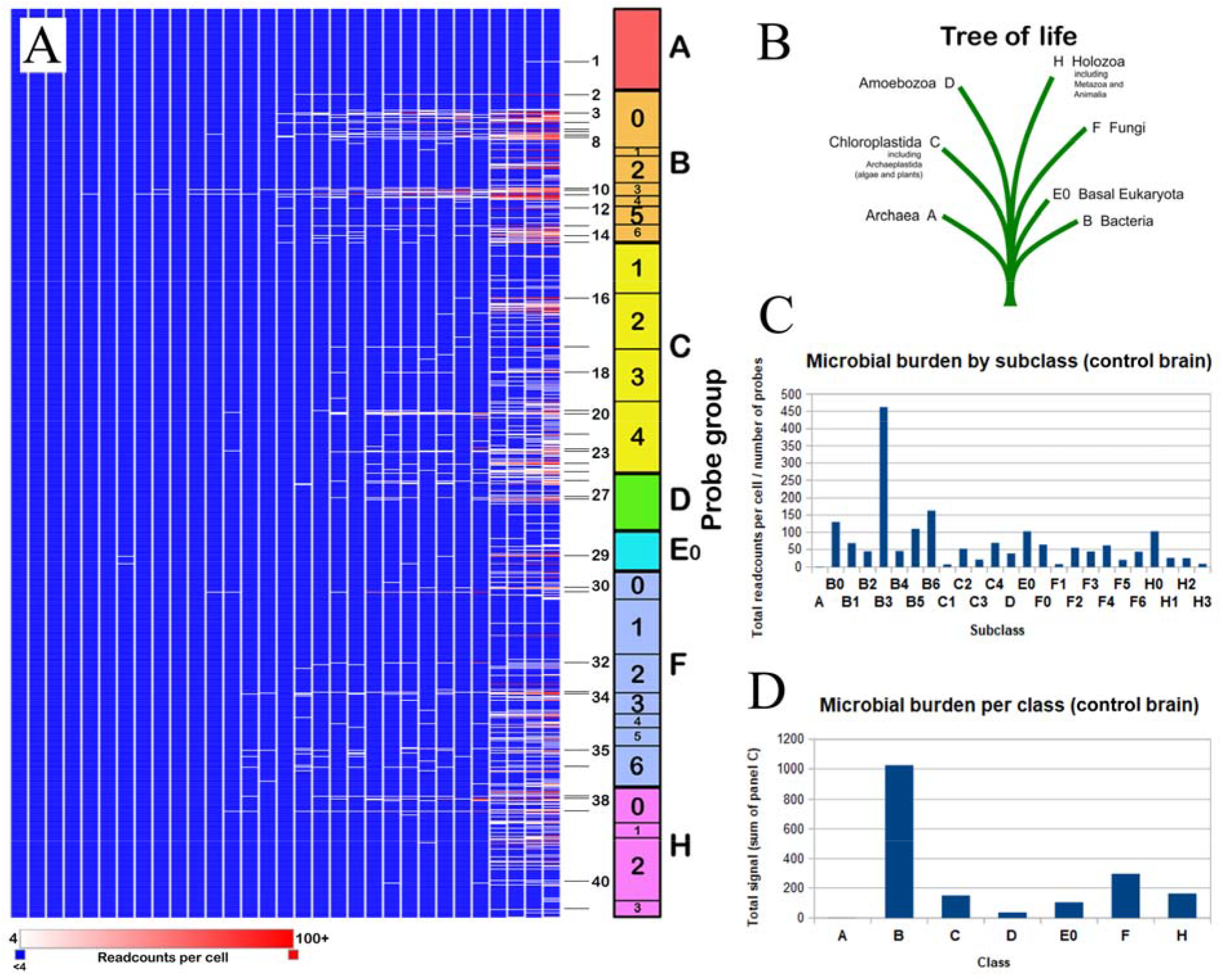
Microbes in normal human brain as assessed by small subunit 16S/18S rRNA sequences (electronic tree of life, eToL (Hu *et al*. 2022)), normalized to endogenous housekeeping genes and calculated as readcounts per host cell. In this and the following figures brain samples are sorted left to right in order of increasing abundance. Signals of interest in this figure are numbered 1–41; formal species identifications confirmed by independent large subunit 23S/28S rRNA sequences are presented in Table S1. Microbial categories are: A, archaea; B, bacteria; C, chloroplastida; D, amoebozoa; E0, basal eukaryota; F, fungi; and H, holozoa/metazoa.

### Conservation of microbial patterns across evolution

To address whether the overall profile of brain microbes is conserved across evolution, we inspected brain RNA-seq datasets from a range of bilateran species ranging from insects, octopus, and vertebrates through to human. Microbes were detected in all cases. The overall profiles were comparable (**Figure 2**). To examine evolutionary relationships, probes for highly abundant signals in bacteria (group B0) and fungi (group F6) were used to retrieve the exact sequences from the original brain RNA-seq libraries. Contigs were assembled, and standard comparison programs (e.g., Clustal Omega) were used to draw phylogenetic trees. As shown in **Figure 2** (right), there was significant conservation in the species identified. Notably, fungal sequences were more highly conserved (the scale for bacteria in Figure 2C is 10% sequence divergence, whereas for fungi in Figure 2B the scale is 1%). We conclude that the evolution of the brain in bilateran species has taken place against a restricted background of fungal and bacterial species.

**Figure 2.**
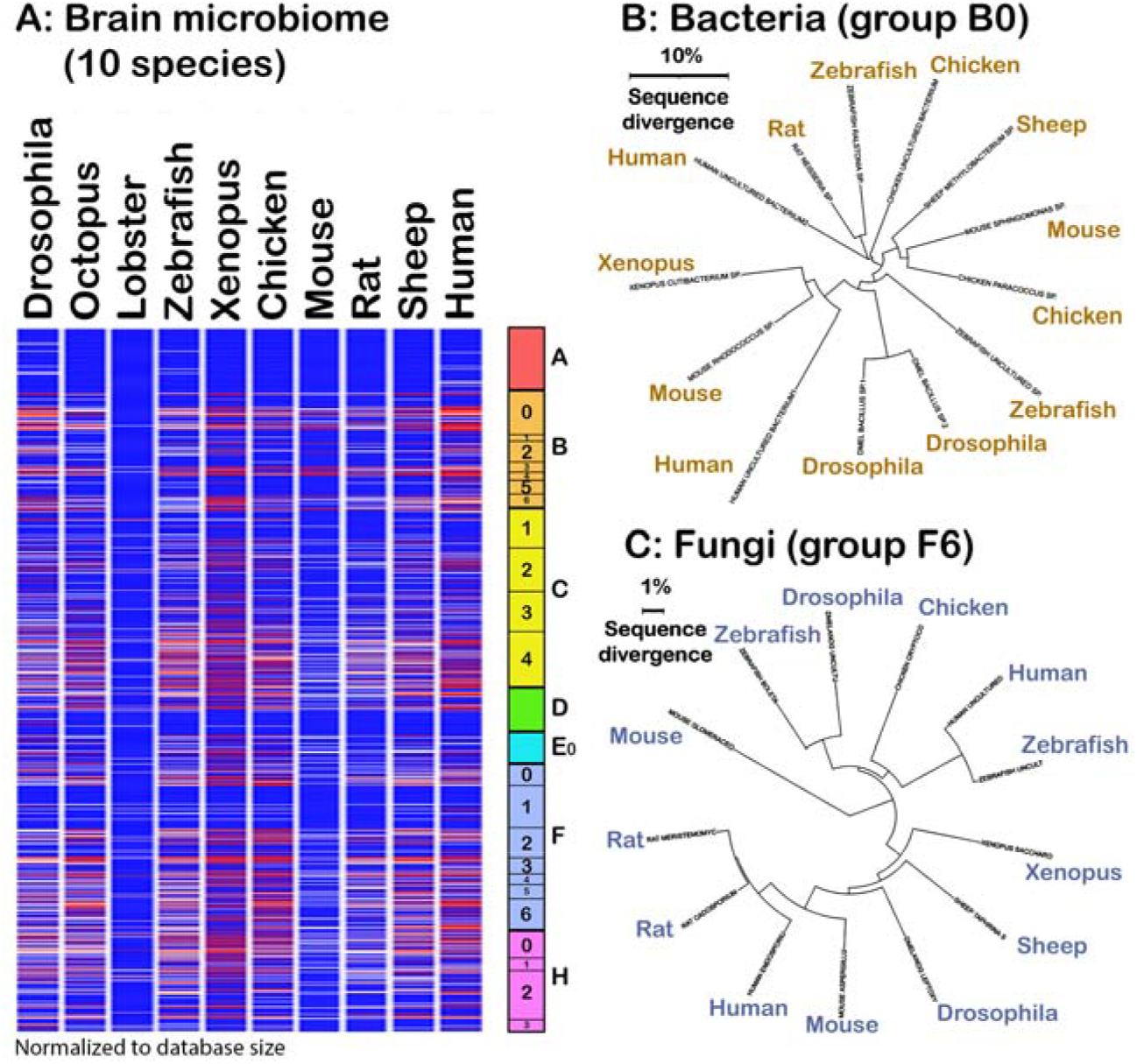
Similar microbial profiles in brains of bilateran species from *Drosophila melanogaster* to human. (A) Microbial profiles across 10 different species. Because normalization to (human) housekeeping gene expression was not possible, intensities were normalized to database size (no scale is therefore presented). In addition, back-subtraction against the host genome/transcriptome was not possible, but all signals inspected by direct sequence retrieval were found to be microbial in origin. (B,C) Sequences matching bacterial group B0 and fungal group F6 were collected from brain samples from different species (octopus and lobster were omitted for simplicity) and used to draw phylogenetic trees based on sequence similarity. Bacterial (B0) sequences were more sequence diverse than for fungi (F6), but there was evidence for evolutionary conservation. For example, the rat bacteria sequence (identified as a *Neisseria*-related species) is 87% identical to the zebrafish sequence (identified as a *Ralstonia*-related sequence) over 1480 nt, although fungal F6 sequences appear to be more conserved: for example 92.52% identity over 1013 nt between the brain F6 fungi of insect (*Drosophila* brain uncultured fungus) and fish (zebrafish brain *Boletacaea* sp.). Abbreviations: ASPERGI LLU, *Aspergillus* sp.; BOLETA, Boletaceae sp.; CRYPTOCO, *Cryptococcus* sp.; DMELANOG, *Drosophila melanogaster*; ENDOSPORIU, *Endosporium* sp.; GLOMERACEO, Glomeraceous sp. LEPTOXY, *Leptoxyphium* sp.; MERISTEMOMYC, *Mristemomyces* sp.; SACCHARO, *Saccharomyces* sp.; UNCULT: uncultured fungus.

### Relationship between the gut and brain microbiomes

We next addressed the relationship between the brain and gut microbiomes. Because gut/fecal sequences are depleted in endogenous human sequences, it was not possible to normalize abundances against housekeeping gene signals. Instead, we evaluated the diversity of different microbes present in the two body regions. Note, these samples are not from the same individuals (we have been unable to source gut/fecal and brain RNA-seq data from single individuals), and future research will need to address this issue.

Few to no species were exclusive to brain; conversely, the majority of signals detected in gut/feces were absent from brain. However, ∼20% of species detected in gut were present in brain (**Figure 3**). These findings suggest that the brain microbiome may be a distinct subset of the microbes present in gut, perhaps indicating that only some microbes have the ability to cross the gut–blood and/or blood–brain barriers. Further studies will be necessary to evaluate the absolute relative abundances of gut (and blood) versus brain microbes.

**Figure 3.**
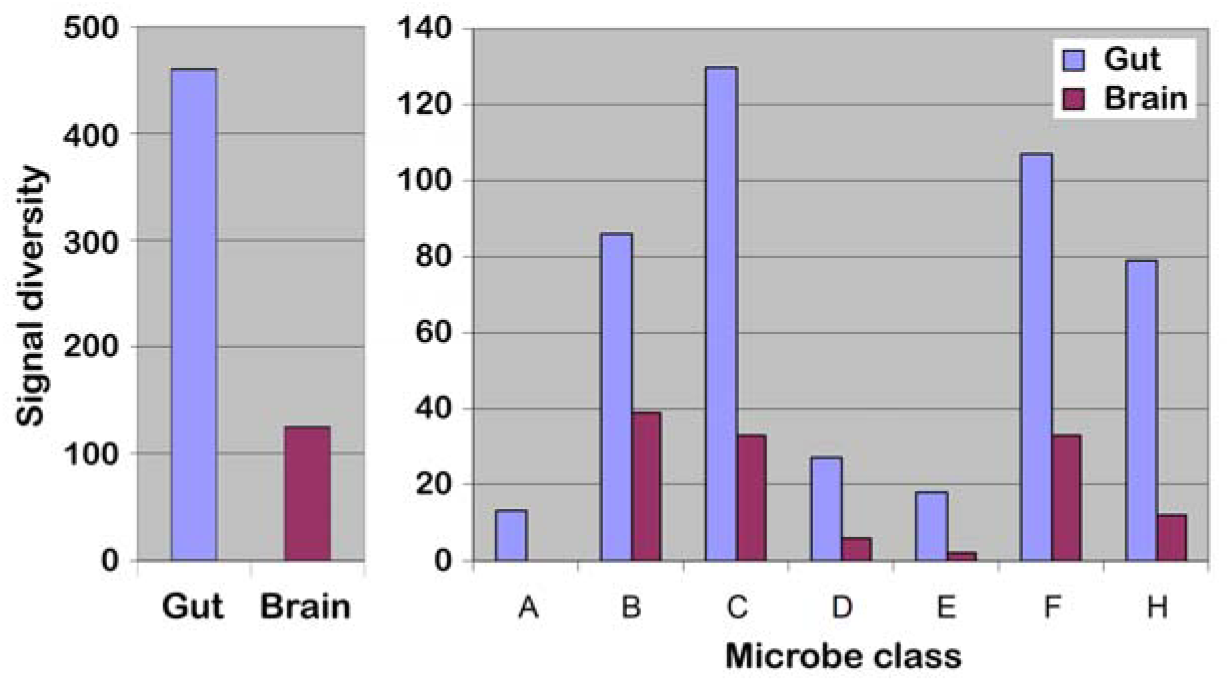
In terms of diversity, the brain microbiome appears to be a subset of the gut microbiome. Microbial groups are classified A–H as in Figure 1. (Left) Total signals in gut and normal brain from 1000+ probes. (Right) Total signals categorized for each microbial group. Because internal normalization to housekeeping genes was not possible for gut samples, absolute abundances could not be determined. Intead, to assess diversity, any signal matching a given probe was considered to be ‘positive’, and the graphs present the total number of probes (signal diversity) in each class that find matches in gut and brain. This demonstrates that the gut microbiome is about fourfold more diverse than the brain microbiome.

### Vulnerable brain regions: focus on limbic brain

Most human brain microbiome analyses to date have been performed on cortex. However, the cortex is not essential for many aspects of cognition, and many individuals with extensive cortical pathology have no cognitive deficits (Discussion). We therefore investigated specific regions of limbic brain where damage is known to be associated cognitive/memory deficits and/or frank dementia including AD. Tissue samples from four different brain regions (amygdala, AMYG; cingulate cortex, BA24; hippocampus, HPC; and hypothalamus, HYPO) from three control brains and six AD brains were obtained from the EBB and subjected to deep sequencing and eToL analysis (35 samples). **Figure 4A** presents a simplified outline of the top 80 signals. Comparative analysis of AD versus control was performed separately (see later below).

**Figure 4.**
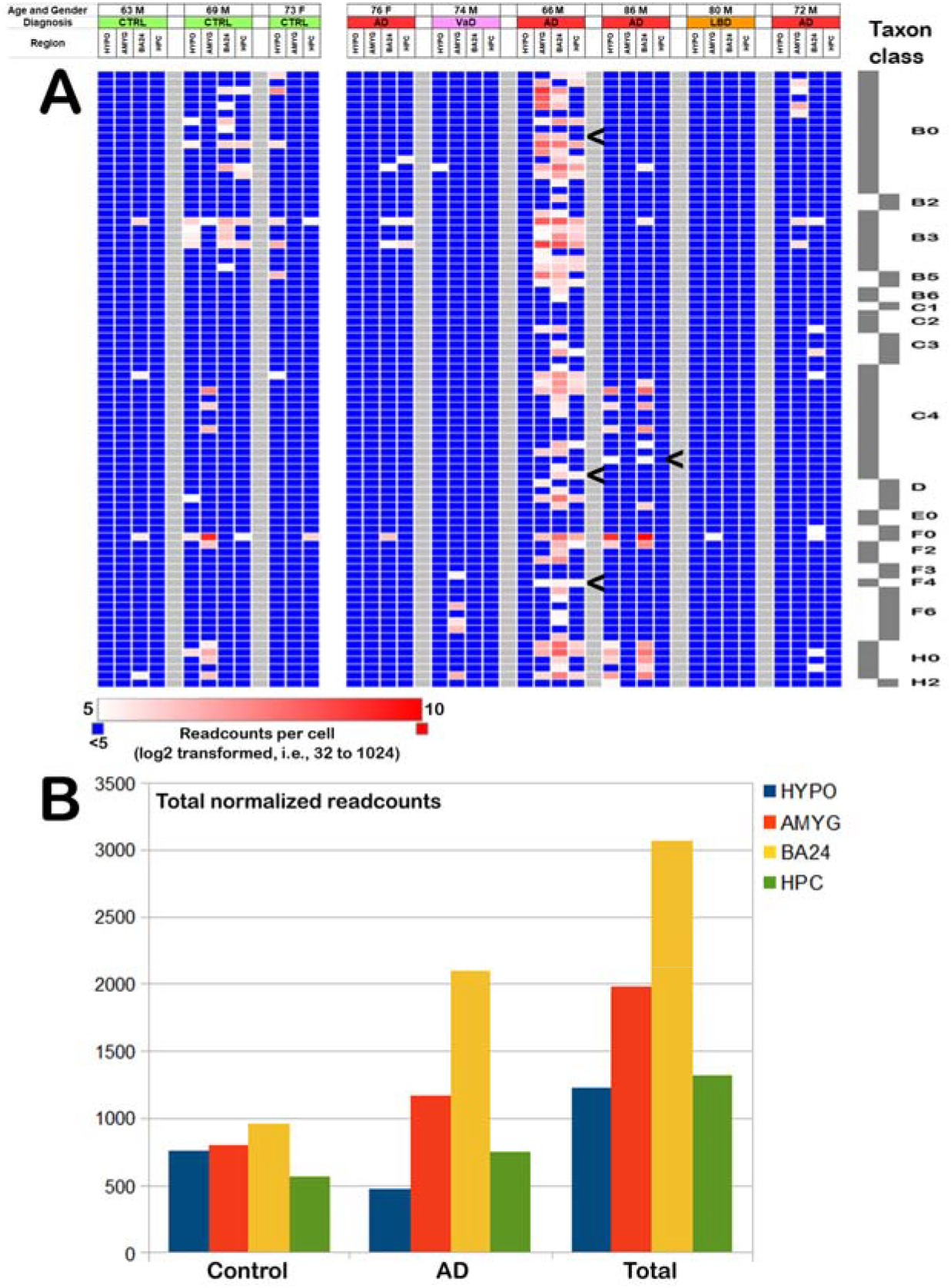
Analysis of Edinburgh Brain Bank (EBB) data. (A) Top 80 microbial signals in terms of abundance. The microbial scale (right) therefore differs from that in other figures. The cutoff has been set highere than in other figures to highlight the detail. Atypical microbes (not in other individuals) present in two brain regions of the same individual are arrowed. (B) Microbes per brain region: highest levels in cingulate cortex (BA24) overlying the hippocampus, and higher levels in cingulate cortex of patients with AD. Because of the small number of individuals sampled, statistical analysis was not undertaken.

The site of microbial infection was found to differ between individuals, irrespective of disease status. For example, one individual had prominent (as defined by our cut-off) levels of microbes only in AMYG, and one only in BA24. In other individuals microbes were restricted to HYPO and AMYG, to HYPO plus BA24, or to BA24 plus HPC. In one individual microbes were found in all four brain regions studied. In some cases atypical microbes were seen in two different brain regions of the same individual (arrowed in **Figure 4A**), suggestive of spreading between brain regions *in vivo*. Overall, microbial burden was highest in cingulate cortex (**Figure 4B**). Other brain regions were not examined.

We investigated whether neuropathology (amyloid deposits and abormal tau/neurofibrillary tangles, NFTs) correlated with microbial burden. Although there was a trend towards a correlation (not presented), this fell short of statistical significance. However, it is important to note that the frozen EBB brain tissues examined were from one side of brain, whereas the fixed sections for neuropathological analysis were from the contralateral side. Although neurodegeneration in AD is often symmetrical, there are cases (for example in hippocampus) where NFT density is not bilaterally symmetrical (Moossy *et al*. 1988), and this could explain the absence of correlation.

### Microbial burden increases with age

To address how the brain microbial burden evolves with age, RNA-seq datasets (human hippocampus) of different ages (*N* = 29) reported in (Kohen *et al*. 2014) were consulted using the eToL method. Matches were principally for bacteria and yeasts. There were no significant differences between males and females, but there was a statistically significant increase in microbial burden as a function of age (*P* = 0.0202) (**Figure 5**). In this cohort only a proportion of the healthy elderly showed an elevated pathogen burden. To determine whether this is a general feature of primates, RNA-seq data from brain (hippocampus) of rhesus macaque (*Macaca mulatta*) aged 10 and 20 years (sample size *N* = 6 in each case) (Xu *et al*. 2018; Xu *et al*. 2020) were analyzed. These Old World monkeys have a median lifespan of around 25 years, but some can live for 40 years in captivity (Mattison *et al*. 2012). Although there are no exact equivalents, 10 years of age in macaque resembles young adulthood in human, and 20 years of age is roughly equivalent to ‘young elderly’ in human. As shown in Figure 5, there was also a trend towards an increase in microbial burden with age (*P* = 0.0944 in view of the small number of samples available).

**Figure 5.**
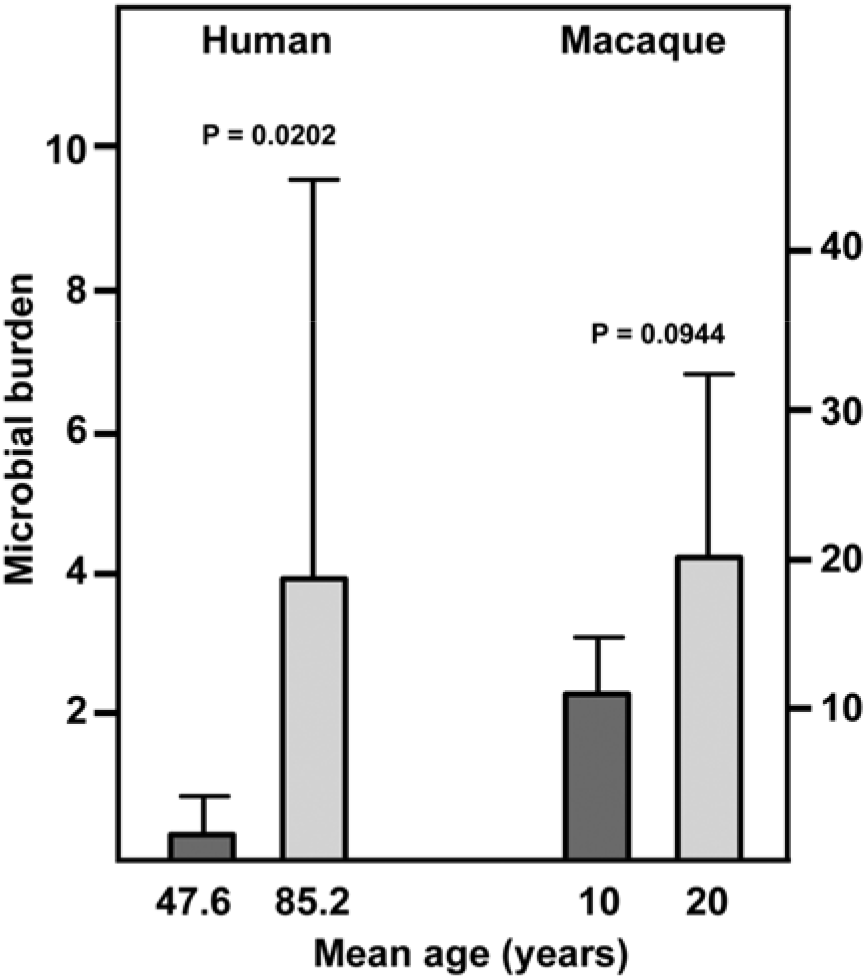
Brain microbial burden as a function of age in (left) human hippocampus (‘young’, *N* = 16, mean age 47.6 years, range 29–59 years; ‘elderly’, *N* = 13, mean age 85.2 years, range 68–95), and (right) macaque hippocampus (‘young’, *N* = 6, all age 10 years; ‘older’, *N* = 6 all age 20 years) (details in **Table S1** and references therein). Error bars are standard deviations, *P* values are from *t*-testing; the non-significance in macaque reflects the small number of samples analyzed. Because different internal controls (housekeeping gene probes) were used for human and macaque, the calculated microbial burdens are not strictly comparable. Note that the expression of the housekeeping genes employed does not change with age. Major species detected in (A) correspond to those reported here and in (Hu *et al*. 2022) (not presented). For macaque, the most abundant microbes were bacteria (Actinobacteria related to *Mycobacterium tuberculosis*, Alphaproteobacteria related to *Rickettsia japonica*, and Firmicutes related to *Bacillus subtilis*; *Bartonella* and *Escherichia* spp. were also seen), fungi (Ascomycota related to *Aspergillus fumigatus*), and apicomplexans (related to *Toxoplasma gondii*). Other species were seen but at lower abundance. No Archaea were recorded in macaque. Viral sequences were not investigated because our methodology only detects high-level homology to human viruses.

### Pooled analysis of AD versus control brain

We next investigated whether any microbes are differentially represented in AD versus control brain. To enhance statistical poer, we pooled all results from the MSBB, EBB, Miami, and Rockefeller studies; this generated a dataset of 31 control samples and 48 AD samples. These were analyzed for differential abundances of cellular microbes, retroelements, and viruses.

#### Cellular microbes

We first performed side-by-side comparisons of the microbial profiles in control and AD brain. As shown in **Figure 6**, the overall profile is conserved across multiple different individuals, although some samples have only low levels of microbes. However, there was no obvious correlation between the overall microbial burden and individual diagnoses of AD versus control (analyzed further below; see also Discussion). Nor was there any evidence at this level of resolution for a specific ‘AD microbe’. The specific identities of these cellular microbes are expanded upon in the next main section.

**Figure 6.**
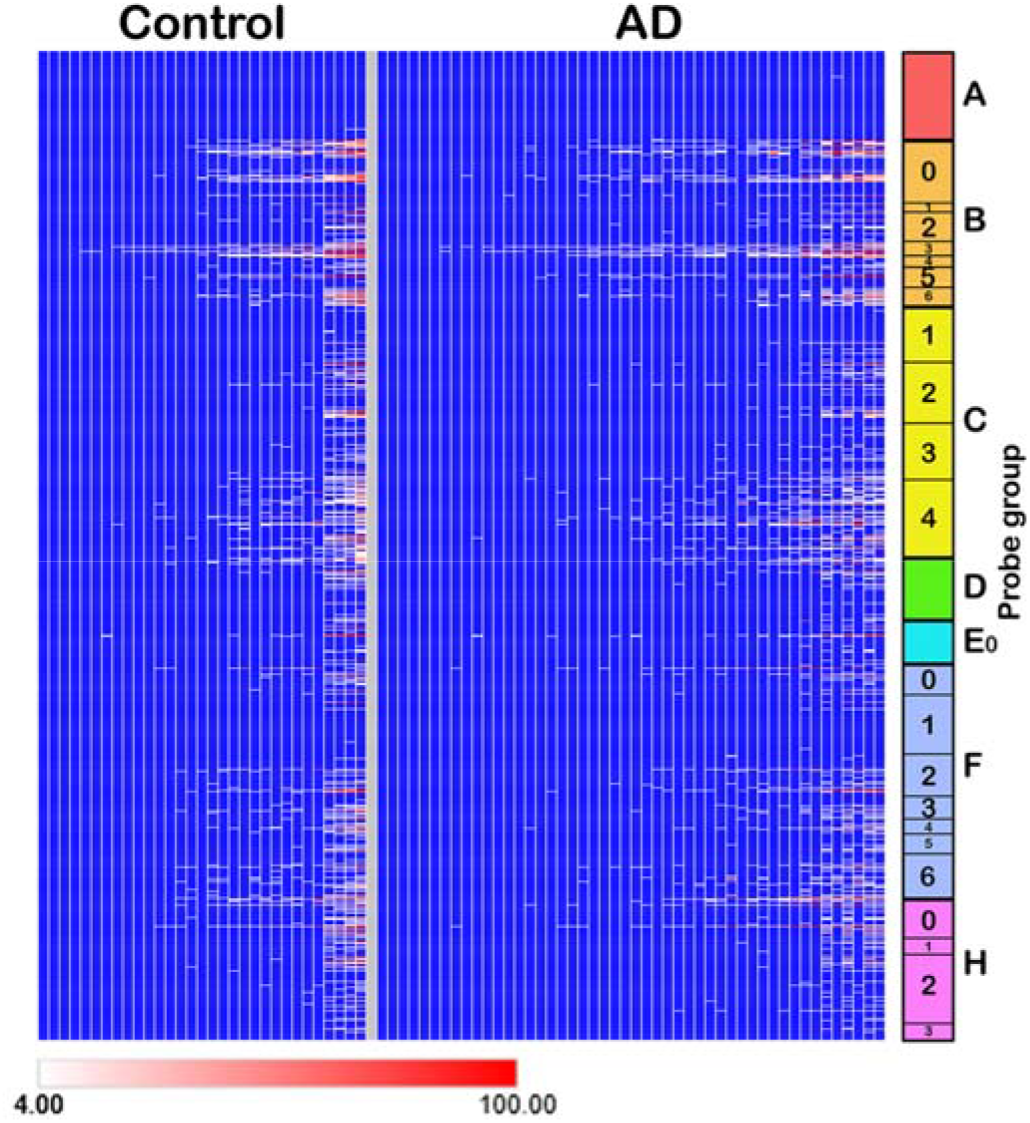
Overall microbial profiles in control and AD brain (arrayed left to right in terms of global abundance) indicating that no specific microbial signals are unique to AD brain. However, this analysis does classifies microbes by group and does not distinguish between innocuous and pathogenic variants within the same group.

#### Viruses and endogenous retroelements

Adenovirus type C (AdC) was the major virus in brain, representing 83% of all viral transcripts across all the datasets, and was detected in ∼50% of all samples (**Figure 7A**). AdC reads were principally principally from the major late transcripts (L1–L3), indicative of productive infection, and to a lesser extent from the E1A/E1B regions (supplementary **Figure S2**). Of note, the AdC substrain detected appeared to be relatively conserved across different individuals ( **Figure S2**). Low-level transcripts of other viruses such as HSV1, EBV, and CMV were found (<0.1 reads per host cell) in some individuals, but the biological significance of this low-level gene expression is not known. Some HHV6A and 6B sequences were detected, but these could not be mapped to the viral transcriptome, and were spread across the entire viral genome (with higher copy numbers in the repeat regions). Our interpretation is that these might represent genomic DNA (e.g., virions), not transcripts. A complexity is that some individuals harbor integrated copies of HHV6A/B (e.g., (Tanaka-Taya *et al*. 2004)), and we have been unable to confirm that the signals observed constitute evidence of biologically relevant infections. There was an inverse relationship in AD brain samples between the presence of HHV sequences and adenovirus C – all AD individuals with detectable HHV sequences were negative for adenovirus C, whereas all individuals positive for adenovirus C were negative for HHVs. However, this was not true for control brain samples (**Figure 7A**, left).

**Figure 7.**
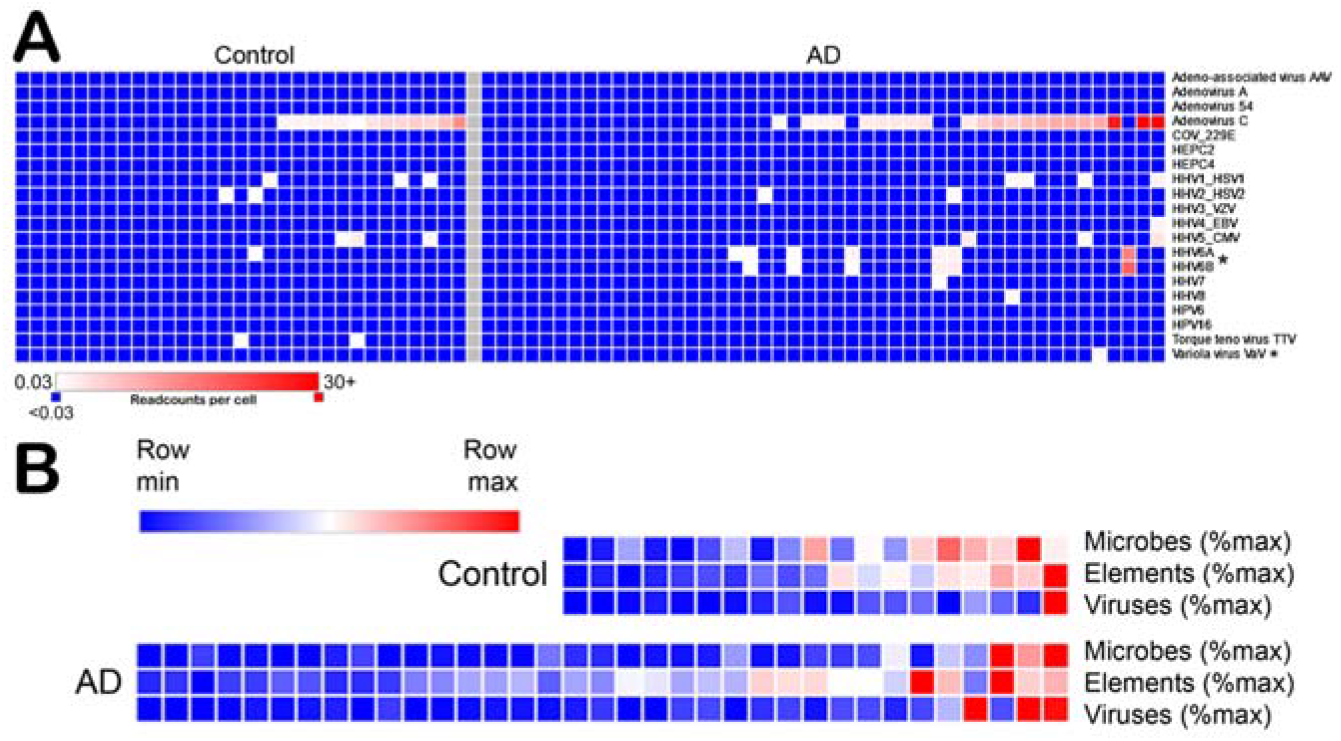
Microbial classes in control and AD brain samples. (A) Viruses: samples are sorted left to right in each case according to the total number of reads (internally normalized); a lower cutoff (0.03 reads per host cell) was employed than for cellular microbes (typically 4–5 reads per host cell) to minimize false negatives. *Text for commentary. (B) Correlation between high abundances of microbes, ‘elements’ (endogenous retroviruses and retroelements), and viruses in both control and AD brain. The coefficients of determination were: cellular microbes versus viruses, *R*^2^ = 0.394; retroelements versus viruses, *R*^2^ = 0.139; and microbes versus retroelements, *R*^2^ = 0.267. Viruses in (A) are HEPC2/4, hepatitis C2/4; HHV1/2, human herpes virus 1/2 (herpes simplex HSV1/HSV2); HHV3, varicella zoster virus (VZV); HHV4, Epstein–Barr virus (EBV); HHV5, cytomegalovirus (CMV); HHV6A/6B/7/8, human herpes viruses 6–8; HPV6/16, human papillomaviruses 6/16; TTV, torque teno virus; VaV, variola major-related virus (vaccinia virus).

Overall, the abundance of viruses in AD brain was somewhat higher than in control brain (*P* = 0.04), although many individuals in both the control and AD datasets had few viral sequences, pointing again to heterogeneity among different cases (**Figure 7A**). This also provided evidence of *in vivo* spreading because the exact substrains differed between individuals, and the same AdC substrain was present in multiple brain regions of a single individual (**Figure S1**).

Retroelements were highly and widely detected in all brain samples, and we also detected transcripts of several endogenous retroviruses; however, there was no clear relationship between the presence of retroelement sequences and disease status (AD versus control; data not presented).

#### Correlation between cellular microbes, retroelements, and viruses

To determine whether the presence of different cellular and viral microbes might correlate with the expression of endogenous retroelements, we took the total number of normalized readcounts for each microbial group in each sample and expressed this as a percentage of the maximum normalized readcount among all samples analyzed. Data were arrayed left to right according to the total signal for all three groups. As shown in **Figure 7B**, there was a correlation where individual samples with the highest abundance of microbes also tended to have the highest abundance of retroelement transcripts and the highest abundance of viral signals (pairwise *R* values between 0.14 and 0.39).

### Over-abundance and over-representation of select microbial species in AD brain

A key issue concerns whether any specific microbial species are more abundant in AD brain than in control brain, noting the complexity that our control brain samples are likely to contain many individuals with pre-AD, and some ‘AD’ individuals are likely to have other conditions, as noted in the EBB cohort (Discussion). We therefore used two different measures of differential representation. First, we inspected the mean numbers of readcounts in AD versus control (i.e., differential abundance). Second, we evaluated the *proportion* (emphasis added) of AD brains that have a higher number of readcounts than control brain (i.e., differential representation). Our conclusions are based on both metrics.

#### Differential abundance

From 16S/18S rRNA readcounts we determined the mean mean abundance level within the control samples in each of the four separate datasets to allow for technical differences in their different methods of preparation. The ratio of the readcounts in the AD samples to the control samples in each dataset was first calculated for all probes, and then averaged across all datasets. Signals were then sorted according to mean relative abundance. This identified a cluster principally of bacteria and fungi that appeared to be overabundant, sometimes by large margins. However, because we used >1000 probes, some differential signals might have been serendipitous (Bonferroni correction). We therefore sought independent confirmation, as described below.

We took the top 100 16S/18S probes that provided evidence of overabundance in AD brain, identified the species by reference to NCBI, and retrieved the corresponding 23S/28S large ribosomal subunit rRNA sequences (in cases where the exact sequence was not available, the closest related species was selected). Using the same protocol as before, random probes were generated, filtered against human sequences, and brain matches retrieved from the two largest datasets – EBB and the selected MSBB data. Readcount abundances versus control were calculated. This confirmed that, for all of these selected probes/species, the mean number of readcounts in AD brain was above that in control brain, sometimes by very large factors (>20; **Figure 8A**). Among the top 100 differentials the mean overabundance was 13.68 (range 2.23 to 181.97; SD 20.40; *P* = 0.01). Because many probes were independently designed from the same species (thus representing a probe ‘cluster’), we determined the mean extent of readcount overabundance for each cluster (individual species represented by multiple probes) (**Figure 8B**). From this cluster analysis the mean overabundance (clusters within the top 100 differentials) was 12.59 in AD versus control (range 2.52–36.88; SD 7.83; *P* <0.001).

**Figure 8.**
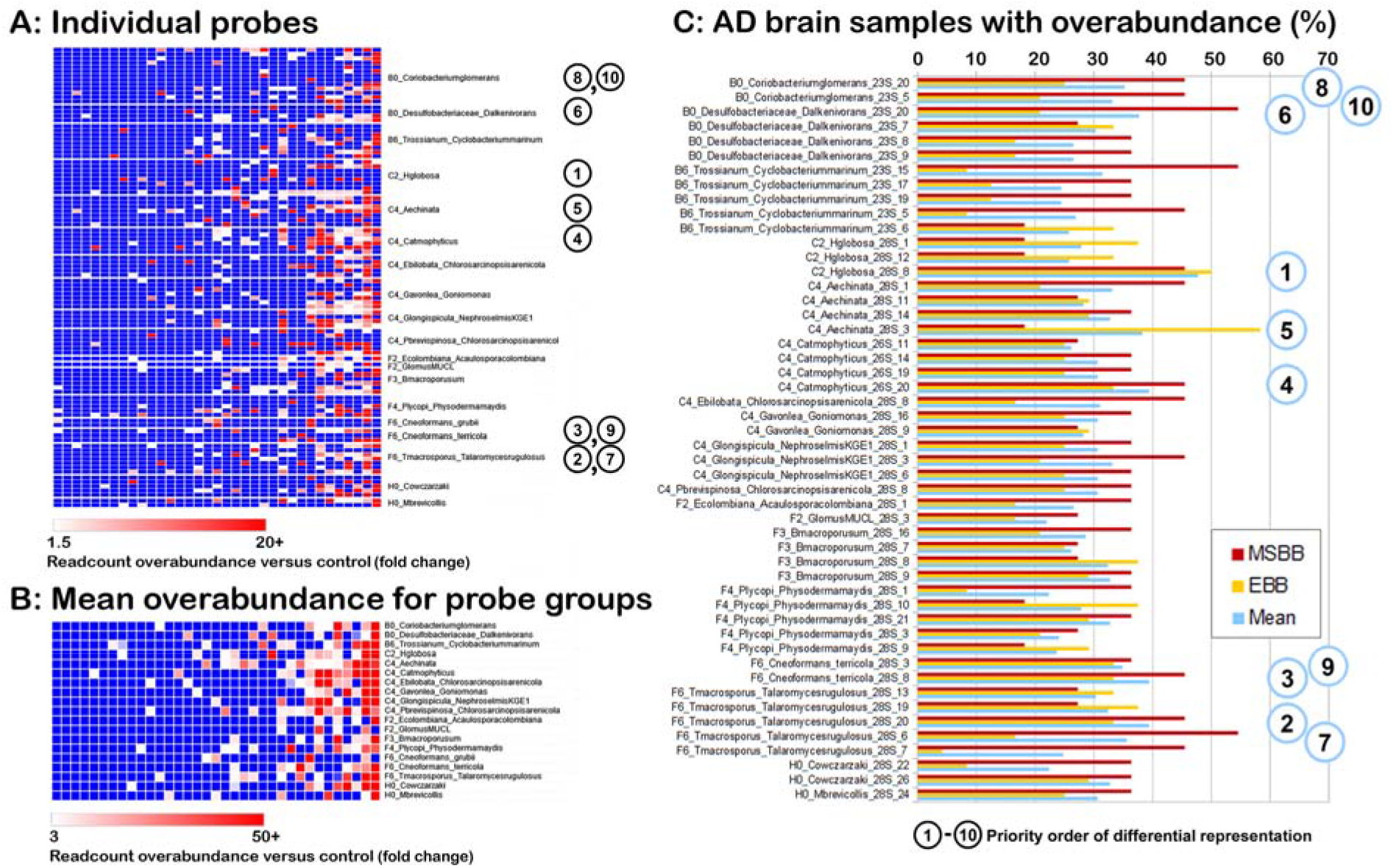
Differential representation in Alzheimer’s disease (AD): species overabundant in AD brain versus control identified from 16S/18S rRNA were confirmed using independent 23S/28S rRNA analysis of the two largest datasets (MSBB and EBB). In panels A and B samples are sorted left to right in terms of overall excess abundance versus control mean. (A) Overabundances in terms of normalized readcounts for select probes per sample for AD brain samples. The top 10 differential signals are numbered according to panel C. (B) Because in panel A multiple probes have been used for each species, the mean overabundances were calculated for each species cluster (noting that these are unlikely to be monophyletic). (C) Over-representation: the proportion of AD brain samples with overabundance. The priority order in terms of mean over-representation (MSBB, red; EBB, yellow; mean of MSBB + EBB, blue) is numbered from 1 (47%, highest proportion of samples overabundant) to 10 (33%, 10th highest in mean differential in the proportion of samples showing overabundance); absolute abundances are presented in Figure 9.

#### Differential representation

We then assessed the second parameter, ‘over-representation’ (the proportion of samples in which there was overabundance of matches in AD versus control). We took the top 50 microbe signals in terms of relative overabundance (previous section), and calculated which signals are most commonly over-represented in AD brain. As shown in **Figure 8C**, increased signals were observed in up to 60% of AD brain samples, although many were only over-represented in a smaller fraction (e.g., 20–40%).

For species identification, to simplify our task we selected the top 10 23S/28S probes in terms of the proportion of AD brain samples displaying overabundance of matching reads (numbered 1–10 in **Figure 8C**). The probes were used to retrieve matches from the most highly affected AD brain samples in the EBB dataset (legend to **Figure 9**), and contigs were assembled. In most cases a single contig was generated that represented the majority of matching reads (if two contigs with similar numbers of read matches were obtained, both are listed). These top contigs were used for species identification by database searching.

**Figure 9.**
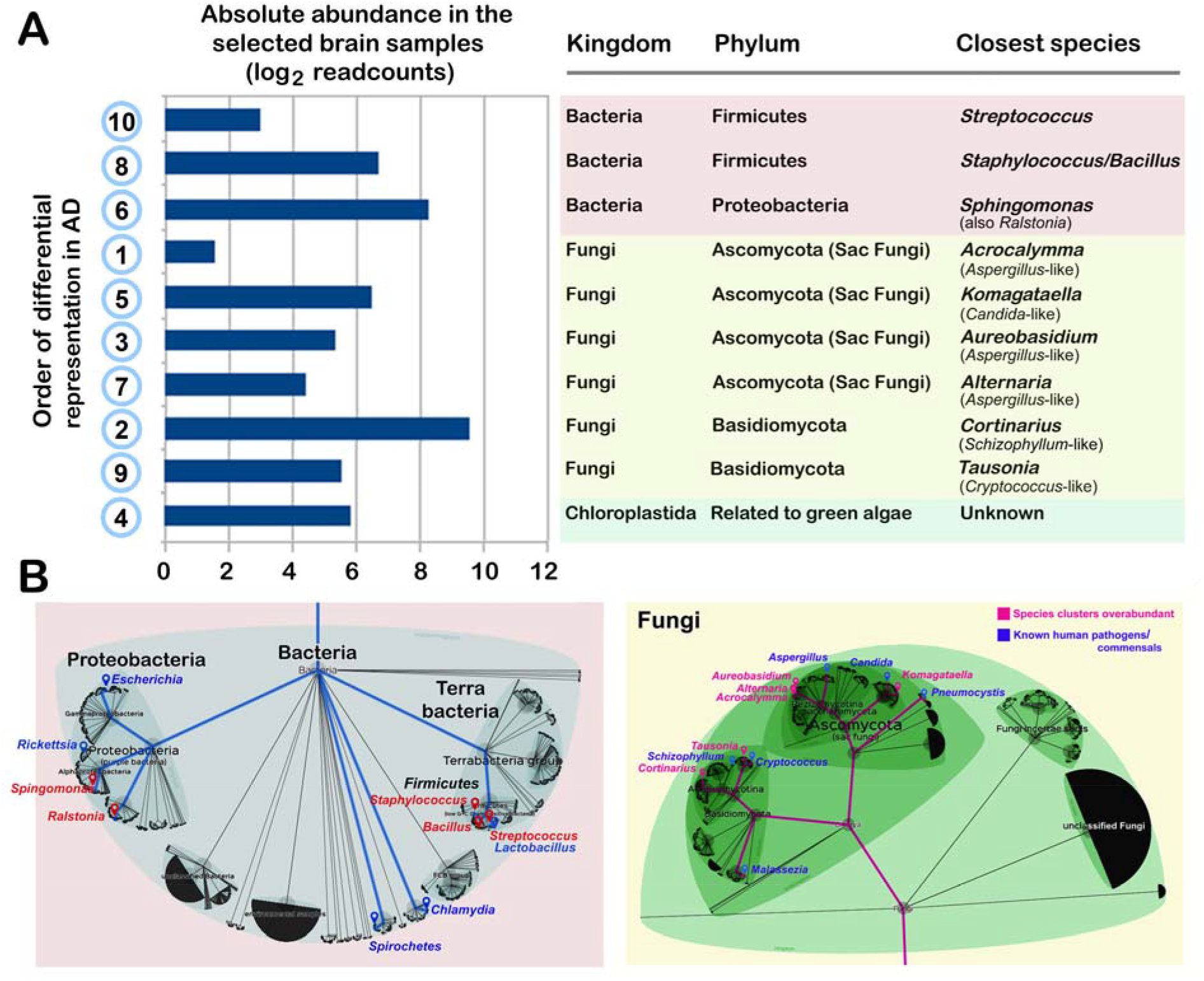
Species over-represented in Alzheimer disease (AD) brain confirmed by 23S/28S rRNA analysis. (A; Left). Priority order 1–10 in terms of differential representation in AD samples versus control samples (proportion of samples showing over-representation), together with the absolute abundance in the AD samples {log_2_ readcounts; ranging from ∼3 (log_2_ = ∼1.5) to ∼1000 (log_2_ = 10)}. (A; Right) Identification of the exact species for the top 10 priority signals (from **Figure 8**) shown on the left. Species identifications are based on all matching sequences from Edinburgh Brain Bank (EBB) brain samples with high-abundance microbial signatures (SD037/18, BA24; SD014/17, HPC; SD014/17, BA24; and SD001/17, BA24; details in **Table S2**). (B) Phylogenies of the different species detected focusing on bacteria (Left) and fungi (Right): typical human pathogens/commensals are shown in blue; the species detected in AD brain are shown in red. The lower panels were drawn based on Lifemap NCBI version at http://lifemap-ncbi.univ-lyon1.fr/.

**Figure 9A** shows the top species overabundant in AD brain versus control, sorted according to microbial class, in terms of both the proportion of AD samples showing differential abundance versus control (left, 1–10) and the absolute abundance in AD brain (log_2_ readcounts). The highest-abundance species over-represented in AD brain versus control (and also the second most differential) was identified as *Cortinarius*, a Basidomycota fungus related to the known human pathogen *Schizophyllum*. Other fungi identified included another Basidomycota, *Tausonia*, that is related to *Cryptococcus*, three *Aspergillus*-like Ascomycota (sac fungi) – *Acrocalymma, Aureobasidium*, and *Alternaria* – and *Komagataella*, that is related to another human pathogen species, *Candida*. Among the bacteria over-represented in AD brain, the Proteobacterium *Sphingomonas* (also *Ralstonia*) was the highest-abundance bacterial species detected, followed by the Firmicutes *Streptococcus* and *Staphylococcus*/*Bacillus*, also known human pathogens.

We previously reported that species identified in human brain are not monophyletic (Hu *et al*. 2022), and instead represent a clade of related sequences. To address this, we used probes corresponding to a representative species, *Malassezia* (a Basidomycota fungus related to *Cortinarius*, and that is present in both control and AD brain, **Table S1**), retrieved all matching sequences from two control brains and two AD brains, and assembled phylogenetic trees. As shown in **Figure S2**, the exact sequences in the four brain samples were not identical. Nevertheless, some subclades did appear to be present in AD brain but not in control brain. However, this is based on a small number of samples; further research will be necessary to determine whether the exact strains differ between AD and control brain.

Overall, our findings based on second-round 23S/28S rRNA analysis are consistent with (and confirm) our observations on AD and control brain based on 16S/18S rRNA, demonstrating that select bacterial and fungal species are over-represented in AD brain, sometimes by a large margin. Further, they confirm that some individuals (and brain regions) have exceptional levels of brain microbes, whereas others appear to have only low levels in the specific brain regions examined.

## DISCUSSION

This is the first study, to our knowledge, that has analyzed the composition of the brain microbiome in health and disease across the entire tree of life, as well as the absolute levels of microbes versus host cells. The central conclusions are as follows. (1) Remarkably diverse cellular species are present in human brain. In addition to fungi and bacteria, we report archaea, chloroplastida, amoebozoa, basal eukaryota, and holozoa/metazoa, taxa that have not been studied previously. We confirm our previous estimate that there are approximately 0.14 bacteria and 0.05 fungi per control host cell (less than 1/10 000 by volume (Hu *et al*. 2022)), although in some seriously affected samples (e.g., AD case male M66 in Figure 4, this rises to 1.8 microbes per host cell). (2) Comparable profiles of brain microbes are present in bilateran species including *Drosophila*, octopus, fish, amphibian, bird, and mammals, and these may thus have accompanied brain evolution in bilaterans, raising the issue of whether some microbes might be beneficial for brain function (Lathe & St Clair 2020). (3) In human, the gut microbiome is far more diverse than that of brain, and brain microbes detected may represent a distinct subset (∼20%) of those in gut, although this remains to be confirmed. (4) Microbial abundance is often restricted to select brain regions, and these regions differ between individuals. (5) Adenovirus type C represents 83% of all viral sequences in brain; the detection of late transcripts is suggestive of ongoing productive infection; few human herpes virus sequences were detected. (6) Select cellular and viral species/subspecies are present in two or more brain regions of the same individual, indicative of *in vivo* spreading.

We provide evidence that particular microbes are over-represented in AD brain. For bacteria, *Streptococcus* and *Staphylococcus*/*Bacillus* species are classical pathogens, whereas *Sphingomonas* has been reported as an emerging pathogen (Ryan & Adley 2010). Among the fungal species, *Tausonia* is a sister species to *Cryptococcus neoformans*, a known human pathogen, *Komatagaella* is a sister species to *Candida albicans*, and *Alternaria, Aureobasidium*, and *Acrocalymma* are closely related to another opportunistic invader, *Aspergillus*. High sequence homologies were detected between AD brain RNAs and these species, for example 98% over 117 nt with *Cryptococcus*, 100% over 106 nt with *Candida*, and 98% over 181 nt with *Aspergillus*. The final species highlighted, *Cortinarius*, is unusually a member of the Agricales subgroup of Basidiomycota, another fungal group where opportunistic infections of human have been reported (Chowdhary *et al*. 2014). An unknown Chloroplastida species was also detected that is possibly related to the green alga, *Nephroselmis* (discussed further below).

We urge caution in the identification of individual species because several groups (bacteria, fungi, and chloroplastida in particular) are extremely diverse, and what we classify as one taxon might in fact be an unusual species from another taxon that has not yet been characterized. In addition, there are concerns that some chloroplastida might reflect lifelong exposure to plant pollens that potentially could enter solid tissues (discussed in (Hu *et al*. 2022)). This also applies to fungi, but the upregulation of chitinases in AD brain (see below) argues against this as being the major explanation, at least for fungi.

A central issue is whether any of these signals might be generated through contamination during sample preparation and/or in the reagents employed (discussed in (de Goffau *et al*. 2018); extended in (Hu *et al*. 2022)). However, brain from germfree mice was reported to be devoid of microbes (Roberts *et al*. 2018), although this preliminary work remains to be confirmed. Ehrlich and colleagues performed a careful study on the brain bacterial microbiome and were able to rule out contamination (Thwe *et al*. 2025). In our work we excluded very low-level (potentially contaminant) signals. Moreover, our brain samples (EBB) were obtained, processed, and sequenced in parallel by the same investigators, but gave 500-fold (or more) differences in readcounts for each probe. In addition, the profiles of all samples were distinct from one another, arguing firmly against the possibility that the signals represent systematic operator or reagent contamination.

Viruses face the same concern. Nevertheless, some sequences detected through homology to variola major virus correspond to vaccinia virus, an attenuated virus vaccine that was widely used in previous generations to eradicate smallpox – smallpox vaccination in the UK and USA was discontinued in 1971/1972, pointing towards long-term persistence of viral sequences in human brain and away from sample/reagent contamination. Luis Carrasco and colleagues have confirmed the presence of bacteria and fungi in AD brain by proteomics, immunohistochemistry, and peptidoglycan analysis, also arguing against contamination (Pisa *et al*. 2015; Pisa *et al*. 2017). In addition to hyphae-like structures, the fungal cell wall component chitin has also been reported in AD brain (Castellani *et al*. 2005; Castellani *et al*. 2007; Pisa *et al*. 2016), but was not found in brain from multiple sclerosis patients (Sotgiu *et al*. 2008). Indeed, upregulation of human endogenous antifungal chitinase mRNA in AD brain (Choi *et al*. 2011; Sanfilippo *et al*. 2016; Pinteac *et al*. 2021; Watabe-Rudolph *et al*. 2012; Magistri *et al*. 2015) is strongly indicative of *in vivo* fungal infection that cannot be explained by sample contamination. In our differential analysis it is implausible that only AD samples (and not control samples) would be contaminated. In sum, the combined weight of evidence argues against the possibility that some of our findings might be ascribed to contamination, although one should remain vigilant for this possibility.

A second issue concerns whether the species we detect are long-term invaders or were recently acquired (*in vivo* biocontamination type 2B in (Hu *et al*. 2022)). For example, the cause of death in AD is predominantly through acute and chronic pulmonary (and perhaps systemic) infections caused by agents such as adenoviruses and associated pulmonary bacteria. Therefore, one might suspect that these could have invaded the brain only well after AD had developed. For fungi, however, Carrasco and colleagues reported that fungal hyphae and glycoproteinaceous structures are present in brain (see above), structures that take months or even years to develop (Alonso *et al*. 2018), arguing against peri-mortem acquired infection of brain tissue, at least for fungi. However, this remains an open issue for other taxa because of the possibility that microbe abundance might increase in the immediate post-mortem period (Emery *et al*. 2022).

On balance, the data argue that diverse microbes are present in AD brain, and some specific groups are overabundant. Could any of these microbes lead to brain damage culminating in AD? Identifying a causal relationship between microbial infection of the brain and AD is not easy, for multiple reasons that have plagued research in the field.

First, control samples are likely to include several individuals with prodromal AD (i.e., normal cognition despite the presence of brain pathology). A key concept in this debate is that of ‘cognitive reserve’ (Stern 2012). The term was introduced with the aim of explaining well-known factors that offer significant protection against AD, such as educational and occupational attainment. Briefly, the concept posits that individuals with better-performing, more flexible, and/or functionally redundant brain circuits can better tolerate damage. This interpretation, now widely accepted, means that – in two individuals with exactly the same extent of brain damage – one may be diagnosed as AD, and the other (with greater cognitive reserve) as ‘cognitively normal’. For this reason we expect that our ‘control’ cohort will contain several individuals with AD-type brain pathology, which would be consistent with our findings, but which is certain to complicate analysis. Conversely, because of diagnostic overlaps between conditions such as vascular dementia, Lewy body dementia, and AD, our ‘AD’ cohort is certain to contain individuals with other non-AD conditions that may or may not share a common pathoetiology.

Second, there is increasing evidence that AD itself is not a singular condition, and has different pathobiological subtypes (Jellinger 2020). Indeed, the EBB cohort of AD patients included one individual with vascular dementia, and one with Lewy body dementia, two conditions that overlap with AD, and both of whom were diagnosed with AD during life. Within AD itself, Zheng and Xu, based on molecular data, classified AD into two subtypes (synaptic and inflammatory) (Zheng & Xu 2021), whereas Levin *et al*., based on positron emission tomography (PET) scanning, classified AD into three different subtypes (Levin *et al*. 2021), of which ‘typical’ (49% of cases) and ‘limbic predominant’ (45%) were the major types. However, it is possible that these share a common pathoetiology, and differ only in the brain regions most affected. A meta-analysis of biological subtypes of AD was recently published (Ferreira *et al*. 2020). I t is not known how these categories correspond to the samples studied in our current work.

Third, the location of damage is likely to be crucial. Studies over the past 30 years have found major discrepancies between overall neuropathology and cognitive impairment: some individuals with marked pathology had no obvious signs of dementia, whereas other subjects with major cognitive impairments had little or no obvious neuropathology. The most intriguing individual in Snowdon’s illuminating Nun Study cohort, Sister Mary, lived to the grand age of 102 without any evident cognitive impairment, but was found postmortem to have more neuropathology (neurofibrillary tangles and neuritic plaques) in her hippocampus than all bar one of the other sisters (*N* = 118) (Snowdon 1997). Conversely, in Fischer’s 1907 paper he remarked that, in four of 16 patients with senile dementia, amyloid plaques were completely absent in the brain regions studied (edited translation in (Bick *et al*. 1987)). Recent work in our laboratory has confirmed this finding, and many non-demented individuals have severe neuropathology, whereas multiple AD patients have little neuropathology in the brain regions examined so far (**Figure S3**).

This argues that even fairly widespread brain pathology *per se* does not produce AD; instead, the specific location of the damage may determine the outcome, as we have argued (Lathe & St Clair 2023), a concept firmly supported by our finding that microbes are abundant in specific brain regions of a single individual, but are largely absent from adjacent regions. Moreover, although we report here the presence of microbes in several brain regions, we have not so far examined other key brain areas. In addition to limbic thalamus (Aggleton *et al*. 2016) and locus ceruleus (LC) where neurodegeneration parallels cognitive decline (Kelly *et al*. 2017), we have been unable to examine brainstem cholinergic nuclei such as the nucleus basilis of Meynert (NBM). This is an important omission because many AD cases respond to medications that increase cholinergic signaling – the cholinergic hypothesis of AD (Bartus *et al*. 1982) – but expert neuropathologists advise that the NBM is almost impossible to dissect in fresh sections because of the lack of histologically discernable features, and future efforts will be necessary to overcome this problem. Other regions warranting attention include the substantia nigra where damage has been implicated in the pathoetiology of Parkinson’s disease, a condition that is sometimes comorbid with AD.

Fourth, we suspect (by analogy to the lung and gut) that there may be a normal brain microbiome, and that some species found in brain may be harmless/innocuous bystanders, or might even be beneficial (Lathe & St Clair 2020). By contrast, some select groups or subgroups could be highly damaging, perhaps by secreting toxins that kill host neurons. Further analysis of toxins will be necessary.

Overall, however, our data point to a pattern of microbial upregulation in around one half of AD patients, and these cases could potentially correspond to one of the AD subtypes described previously (see above). To address this, future work in the field will need to accurately subclassify AD into different subtypes. However, this leaves open the possibility that neuropathology in roughly one half of AD cases is not associated with microbes and has a different pathoetiology (e.g., chemical/environmental). Nevertheless, we have not so far looked at other potentially crucial brain regions in these individuals, and/or explored only one side of the brain; we stress that our ‘non-microbial’ cases of AD may have infections in other brain regions or in the other side of the brain.

Despite these caveats, the fungal microbes we identify are remarkably similar to (and confirm) those previously suggested to be overabundant in AD brain, specifically fungi related to *Cryptococcus/Malassezia, Candida*, and *Aspergillus* (Alonso *et al*. 2014; Pisa *et al*. 2015; Pisa *et al*. 2017; Alonso *et al*. 2018). Confirmed brain infections with *Cryptococcus* have previously been diagnosed as dementia/AD (e.g., (Steiner *et al*. 1984; Ala *et al*. 2004)). The bacterial species identified are also similar to those previously reported (e.g., (Emery *et al*. 2017)), noting that bacterial infection of the brain can also cause dementia (e.g., (Gonzalez *et al*. 2019)). In general, the species we report are known causes of human pathology. This includes viruses. The finding that adenovirus is the major virus present in human brain, both in controls and in AD, is of interest because adenovirus is a known cause of brain disease including meningitis in immunocompetent children (Schwartz *et al*. 2019), although more rarely in adults. In addition, adenoviruses can cause long-term CNS inflammation in animal models (Dewey *et al*. 1999).

In AD, of all the differential signals, fungi related to *Schizophyllum, Aspergillus, Tausonia*, and *Candida* are the most prominent, whereas *Sphingomonas* and *Staphylococcus* and related species are the predominant bacterial differentials. Given that fungi have more ribosomes than bacteria (Hu *et al*. 2022), normalization for this factor suggests that they many be present in brain in approximately equal cell numbers. Fungi and bacteria often synergize for growth both *in vitro* and in a clinical setting (Förster *et al*. 2016; Deveau *et al*. 2018; Little *et al*. 2021), and AD may therefore be associated with synergistic polymicrobial infections, as previously speculated (Pisa *et al*. 2017).

These findings suggest that AD may be a polymicrobial disease in the same sense that pulmonary disease is polymicrobial – pneumonia-like symptoms can be caused by influenza virus in one individual, by coronavirus in a second, and by bacteria such as *Streptococcus pneumoniae* or *Haemophilus influenzae* in another, or even by fungi such as *Aspergillus fumigatus* in others. These pathogens often contribute synergistically to pulmonary disease.

The finding of an uncharacterized chloroplastida (algae/plant-related) species in AD brain is of interest because cases of human infection with algae have been reported (Lass-Flörl & Mayr 2007) and an algal virus of unknown provenance found in human samples was associated with cognitive impairment (Yolken *et al*. 2014). *Toxoplasma* spp. (the etiologic agents of toxoplasmosis) are also broadly classified as chloroplastida and have previously been implicated in brain disease including schizophrenia and dementia (Yolken *et al*. 2009; Yang *et al*. 2021). Further research will be necessary to characterize the chloroplastida species detected in AD brain and evaluate their potential contribution to pathology.

Despite these suggestive findings, there is a need for independent confirmation using techniques that are not based on nucleic acid detection, such as direct culture of live microbes from brain tissue, and work in this direction is ongoing (D. Corry, personal communication).

Looking wider, our analysis fits with theories of programmed aging (Goldsmith 2013; Skulachev & Skulachev 2018) leading to immune system decline (immunosenescence (Makinodan 1980); (Lathe & St Clair 2023; Lathe & St Clair 2026) for extensive review). Indeed, the cause of death in AD is not from loss of normal cognition but is typically from pulmonary infection (e.g., (Kukull *et al*. 1994; Beard *et al*. 1996) and/or atherosclerosis and cardiovascular disease (also associated with infection (Dunn 2004; Lathe *et al*. 2014)). In this context it would be of interest to determine whether the microbes found in AD brain are a specific subset of (and thus represent migration from) the microbiome of the gut (or of other tissues), or have an independent origin (perhaps being acquired through the olfactory system).

In conclusion, we have identified a shortlist of microbes that are overabundant/over-represented in AD brain and which may contribute to neurodegeneration. Our results confirm and extend previous reports of polymicrobial infections of AD brain, which need to be corroborated by culture of microbes from AD brain tissue and/or onwards transmission to rodents ((Branton *et al*. 2013); David Corry, personal communication).

However, association does not prove causation, and one must keep an open mind about whether microbes are the cause of neuroinflammation and neurodegeneration in AD, or whether they merely invade tissue that is already degenerating from another cause. Further analysis in animal models will be necessary to evaluate whether specific microbes are, or are not, causally implicated in the pathoetiology of AD. However, a series of case reports have emerged where antimicrobial intervention sometimes brought remarkable benefits to patients with dementia (Lathe *et al*. 2023), and antimicrobial medications offer a potential way forward, noting that the exact treatment may need to be tailored to the exact species present (which would depend on the individual), and would therefore require the development of *in vivo* detection techniques (Lathe *et al*. 2023).

Boosting innate immunity by administration of vaccines and/or adjuvants may also be protective (Greenblatt & Lathe 2024a). For example, the risk of developing AD was 4.8-fold lower in bladder cancer patients receiving BCG vaccine than in untreated individuals (Gofrit *et al*. 2019), and smaller but significant reductions have been seen with other vaccines (Greenblatt & Lathe 2024b). These and other strategies to enhance the immune system warrant investigation (Vaccines for Dementia Study Group 2026). In addition, given the abundance of *Aspergillus*- and *Cryptococcus*-like microbes in AD brain, and their involvement in other human diseases, research to develop vaccines against these and other fungal groups may be warranted (Levitz 2017; Caballero Van Dyke & Wormley, Jr. 2018; Oliveira *et al*. 2021).

In sum, we provide evidence that the brain harbors its own remarkably complex microbiome that extends beyond bacteria and fungi to include chloroplastida and other taxa, some of which are overabundant/over-represented in AD. Future analyses will need to extend microbial analysis to include other brain regions, and similar analyses are warranted in other disorders such as Parkinson’s disease, schizophrenia, multiple sclerosis, and atherosclerosis where an infectious component has long been suspected.

## ACKNOWLEDGMENTS

We thank Ben Readhead (Arizona) for his help in reconstructing datasets, NCBI (Bethesda) for enabling online database searching, and Heiko Braak (Ulm) for help in prioritizing brain regions for analysis. We thank David Breen (Edinburgh) and David Corry (Houston) for helpful comments on the manuscript, and Alasdair Ivens (Edinburgh) for bioinformatic assistance. Alison Daniels (Edinburgh) is thanked for her valuable contributions to this project. This work was funded in part by the Benter Foundation (grant VIRADE to J.H. and R.L.).

## Notes

### Competing Interest Statement

The authors have declared no competing interest.

### Summary of Updates

This paper was filed in 2023. We have made only minor changes to the manuscript to bring it up to date. Our intention is to submit for mainstream publication via bioRxiv.

